# Serial position effects reflect the integrity of distinct brain regions in Major and Minor Neurocognitive Disorder

**DOI:** 10.1101/662742

**Authors:** Nancy S. Foldi, Kirsten I. Taylor, Andreas U. Monsch, Marc Sollberger, Sasa L. Kivisaari

## Abstract

Serial Position Effects (SPE) in wordlist learning provide a rich set of metrics of cognitive functioning. In this study, we systematically mapped the neuroanatomical correlates of SPEs across delays in Major and Minor Neurocognitive disorders. Primacy, middle, and recency SPE scores of the California Verbal Learning Test at Learning, Short delay, and Long delay in healthy controls and patients with neurodegenerative disease were correlated with MRI gray matter signal intensities (voxel-based morphometry, VBM). The VBM analyses revealed distinct patterns of brain-behavioral correlations depending on both the SPE and the time-point. Unlike patients, the healthy controls’ performance incrementally improved recall of primacy items from Learning to Short and to Long delay, i.e., primacy progression. Moreover, the proportion of correct primacy items recalled at long delay compared to Learning correlated with bilateral medial temporal lobe regions, which commonly bear the brunt of pathology of Alzheimer’s disease. The results implicate the primacy progression score as an accessible and sensitive measure for disease detection, progression, and therapeutic response in Major and Minor Neurocognitive disorders.

## Introduction

Words positioned at the beginning (i.e. primacy) and end (recency) of a word list are better recalled than words positioned in the middle of the list. These serial position effects (SPE)^1, 2^ are behaviorally well-documented and putatively reflect distinct underlying cognitive processes^3^. Specifically, primacy item learning is thought to depend more heavily on semantic memory processes than recency item learning, which is posited to rely disproportionately on auditory rehearsal^4, 5^. This account leads to the prediction that primacy and recency recall are disproportionately affected by damage in the medial temporal lobe (MTL) and lateral temporal lobe, respectively, as these brain regions have been shown to differentially contribute to semantic memory and auditory rehearsal processes^6–10^. In the current voxel-based morphometry (VBM) study, we test this hypothesis in the context of Major and Minor Neurocognitive disorder. We systematically map the neuroanatomical correlates of each SPE across Learning, Short delay (SD) and Long delay (LD) recall. We predict that poorer primacy item recall is associated with decrease in gray matter volume in the MTL and suggest that this measure may serve as a sensitive marker of AD-related pathology.

Behavioral assessments of SPE at learning are well studied in healthy adults and persons with Alzheimer’s disease (AD). Healthy individuals show a classic ‘U-shaped’ profile with comparable numbers of primacy and recency words, but fewer from the middle position. We use the term ‘J-shaped’^11^ profile to describe fewer primacy than recency items at learning, which is a well-documented characteristic performance in patients with diagnosed with AD^11–15^ as well as in healthy individuals at risk for AD^16–18^. The model’s divergent patterns ascribe primacy item recall to consolidation of ‘deeper’ semantic elaboration processes, and recency item learning to ‘shallow’ online phonological rehearsal^19^, which is relatively preserved during early stages of most forms of dementia^20^.

Distinct cognitive subprocesses serving primacy and recency item recall at learning lead to the hypothesis that they are subserved by differential neuroanatomical bases. Namely, primacy item processing would be expected to rely on regions associated with semantic processing, such as middle (MTG) and inferior temporal gyri (ITG)^10^ and medial temporal lobe (MTL) structures, which support semantic memory^8^. In contrast, recency items would likely rely at least on right STG which has been linked with phonological processing^7 19^. This anatomical differentiation has received some support in the literature. For instance, reduced primacy item learning in patients with MCI was shown to be associated with reduced gray matter intensity in e.g., temporal pole and hippocampus^15^.

Delayed SPE measures have been less studied, but carry a particular significance in Major and Minor Neurocognitive disorder, given that impaired episodic memory consolidation is the hallmark of Alzheimer’s disease^21, 22^. Although previous studies have found associations between primacy recall at delay and hippocampal volume^16, 23^ and cognitive decline^24^, we claim more targeted steps need to be taken to grasp the full potential of the delayed primacy recall measure. First, we suggest it is important to consider the difference between performance at initial learning and subsequent delayed recalls to tap the individual rate of forgetting or of the ability to conserve primacy items over time. Second, it is important to consider brain regions beyond the hippocampus since the neighboring areas, in particular the perirhinal cortex, have been shown to carry an important role in semantic processing and item-specific memory^6, 25–27^.

The goals of the present study were to systematically determine the functional neuroanatomy of SPE at Learning, at SD, and at LD recall in healthy individuals and patients with Major and Minor Neurocognitive disorders. We anticipated that each group would show the respective U-shaped or J-shaped behavioral profiles reflecting preserved or impaired primacy item learning^14, 28^. Over time at SD and LD, only the patient group would demonstrate a drop in recency items^29^, whereas controls would be able to maintain the initial learned primacy items. Our first hypothesis proposed that brain-behavior correlation analyses would demonstrate a reliance of primacy item learning on MTG, ITG, and MTL^8, 10^ and recency item learning on the superior temporal gyrus (STG)^7, 19, 30, 31 9^. Second, we hypothesized that the range of decay or improvement of primacy items would be associated with volume in the MTL encompassing the hippocampus, entorhinal and perirhinal cortices. These are regions key regions in the early diagnosis of AD as they are affected very early in the course of AD^32^.

## Results

### Behavioral results

Analyses revealed the expected HC over PAT superiority in performance overall (F(1,127) = 193.4, *p* < .001; partial eta squared (*η*^2^) = .60). We found a significant main effect of List Position (F(2,254) = 3.4, *p* = .04; *η*^2^ = 0.03) with better primacy item accuracy than either middle (*p* < .001) or recency items (*p* <.001), which in turn did not differ. There also was a significant main effect of Time (F(2,254) = 3.4, *p* = .03; *η*^2^ = .03) with significant difference between Learning and SD (*p* < .05) and LD (*p* <.005), which were not significant from one another. While age was a significant covariate (*p* = .002) and interacted with Time (*p* = .03), it did not significantly interact with SPE. The important three-way interaction was significant (F(4,508) = 6.6, *p* < .001, *η*^2^ = 0.05; (Fig. 1)). At Learning: each group showed their predicted U-shaped and J-shaped learning profiles (HC: primacy = recency; PAT: primacy < recency, *p* < .001). Patients performed worse than controls on primacy and middle items (both *p* < .001), and a trend for worse performance on recency items (*p* = .08). After interference, both groups’ SPE profiles change from quadratic functions at Learning to linear functions at SD and LD, where patients recalled overall fewer items than controls at all list positions (*p* < .001). Only patients, but not controls, showed the significant drop in recency item recall from Learning to SD (*p* < .001) and LD (*p* < .001), which in turn did not differ (n.s.). Importantly, the controls’ primacy item performance improved significantly over time from Learning to LD (*p* < .001), while numerically improving stepwise from learning to SD (*p* = .05, trend.) and between SD to LD (n.s.). In the patient group, primacy recall remained at a level unchanged from Learning to either SD or LD (n.s.), and showed no improvement. Recency items in the patient group showed a clear decrement from Learning to SD (*p* < .001) and from Learning LD (*p* < .001), while there was no difference between the delays (SD – LD, n.s.). In contrast, the control group maintained recency item accuracy from learning to SD and to LD (n.s.).

**Figure 1.**
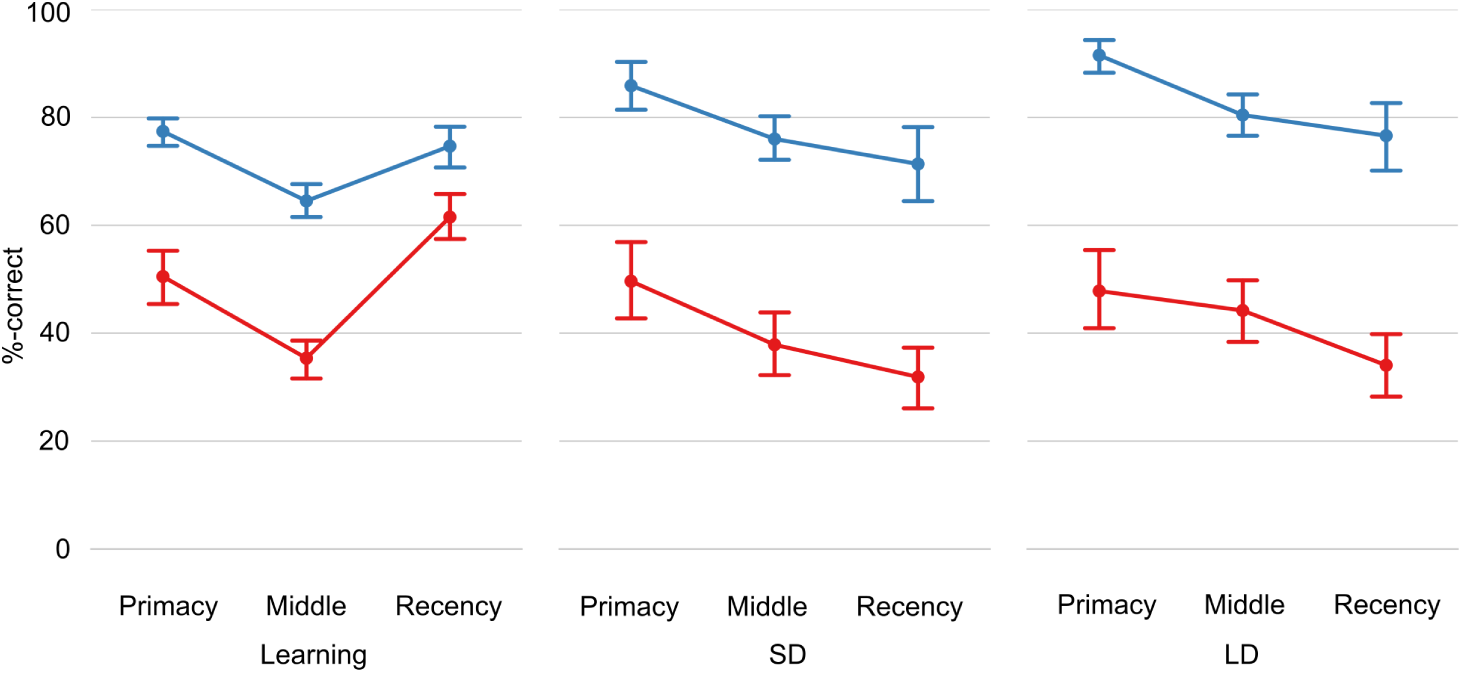
Performance accuracy for primacy, middle and recency serial list positions at Learning, Short Delay (SD) and Long Delay (LD) on the California Verbal Learning Test. Plots indicate mean percent correct for Healthy Controls (blue lines) and Patients (red lines). Error bars indicate 95%-confidence intervals computed using bootstrapping.

### Voxel-Based Morphometry Results

#### Learning

*Primacy*: Poor primacy item performance was associated with reduced GM volume in two clusters (Fig. 2a). One was centered in the right angular gyrus and extended to the right middle occipital gyrus. A second cluster was centered in the left MTG and included the left temporal pole, MTG/STG, hippocampus and parahippocampal gyrus. *Middle*: Poor middle item learning was associated with reduced volume in the bilateral cerebellum (see Table 1). *Recency*: Poor recency item learning was associated with reduced volume in two clusters, one centered in the left hippocampus extending to the parahippocampal cortex and amygdala. The second cluster was centered in the right superior temporal pole extending into the right parahippocampal cortex, MTG and STG (see Fig. 3). The VBM results are summarized in Table 1.

**Table 1.** VBM result summary. Overview of the VBM results demonstrating the cluster- and peak-level statistics for the three effects of interest (primacy, middle and recency) in the different VBM analyses (a) Learning, (b) Short Delay and (c) Long Delay and (d) the difference scores (Long Delay— Learning).

|  | cluster-level<br>(FDR) |  | peak-level<br>(uncorrected) |  | coordinates |  |  |  |
| --- | --- | --- | --- | --- | --- | --- | --- | --- |
|  | p | k | p | t | x | y | z | AAL |
| **(a) Learning** |  |  |  |  |  |  |  |  |
| Primacy | .0001 | 6008 | .0001 | 4.2 | 51 | -64 | 37 | Angular R |
|  | .0001 | 5348 | .0001 | 4.0 | -48 | -27 | -8 | Temporal Mid L |
| Middle | .0001 | 25220 | .0001 | 5.0 | 28 | -40 | -51 | Cerebellum R |
| Recency | .044 | 2008 | .0001 | 4.1 | -20 | -10 | -18 | Hippocampus L |
|  | .044 | 1792 | .0001 | 3.5 | 46 | 10 | -18 | Temporal Pole Sup R |
| **(b) Short delay** |  |  |  |  |  |  |  |  |
| Primacy | .002 | 3928 | .0001 | 4.4 | -48 | 2 | -35 | Temporal Inf L |
|  | .034 | 1986 | .0001 | 3.7 | 52 | -78 | -5 | Occipital Inf R |
|  | .045 | 1685 | .0001 | 3.6 | -14 | -1 | -27 | Parahippocampal L |
| Middle | .001 | 3922 | .0001 | 3.5 | -6 | -69 | -44 | Cerebellum 8 L |
| Recency | .005 | 3201 | .0001 | 3.8 | 36 | -45 | -54 | Cerebellum 8 R |
|  | .036 | 2209 | .0001 | 3.4 | -30 | -42 | -53 | Cerebellum 8 L |
| **(c) Long delay** |  |  |  |  |  |  |  |  |
| Primacy | .0001 | 5471 | .0001 | 4.4 | -30 | -4 | -20 | Hippocampus L |
|  | .001 | 4373 | .0001 | 4.1 | 27 | -7 | -18 | Hippocampus R |
| Middle | ns. |  |  |  |  |  |  |  |
| Recency | ns. |  |  |  |  |  |  |  |
| **(d) Long delay vs. Learning** |  |  |  |  |  |  |  |  |
| Primacy | .0001 | 5597 | .0001 | 4.34 | 30 | -10 | -36 | Fusiform R |
|  | .003 | 3021 | .0001 | 4.2 | -33 | -1 | -26 | Parahippocampal L |
| Middle | ns. |  |  |  |  |  |  |  |
| Recency | ns. |  |  |  |  |  |  |  |

**Figure 2.**
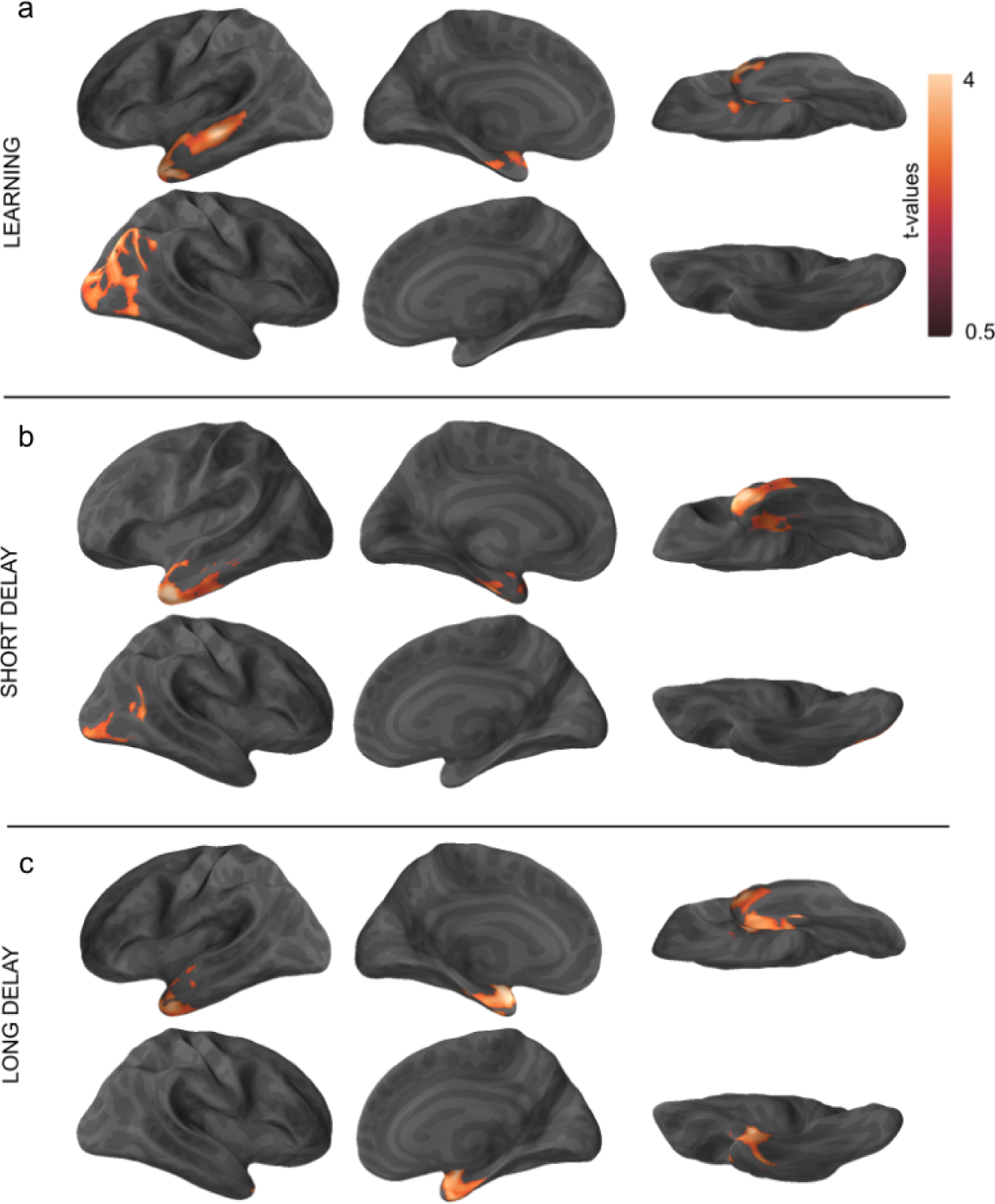
Brain regions where reduced signal intensity significantly correlated with poor accuracy of primacy at a) Learning, b) SD and c) LD. The *t*-values of voxels in significant clusters are project onto the cortical surface.

**Figure 3.**
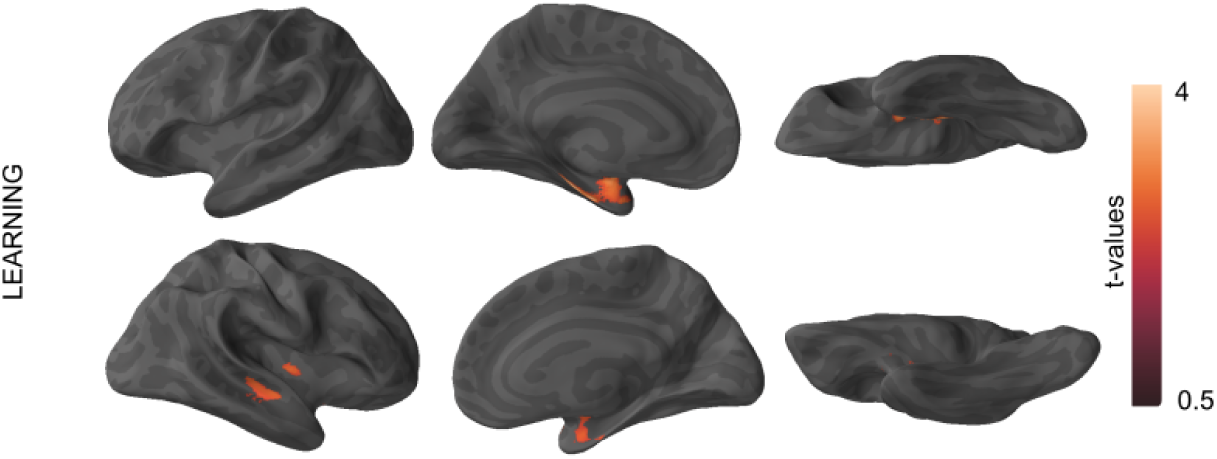
Brain regions where reduced signal intensity significantly correlated with poor accuracy of recency item learning. The t-values of voxels in significant clusters are projected onto the cortical surface.

#### Short Delay

*Primacy*: Poor primacy item recall at SD was associated with reduced volumes in three clusters. One cluster was centered in the left ITG extending into the MTG and temporal pole area. A second cluster encompassed the left MTL region including the hippocampus and parahippocampal gyrus. A third cluster was centered in the right occipital inferior gyrus and extended in the right MTG and angular gyrus (Fig. 2b, Table 1). *Middle*: Poor middle item recall at SD was associated with reduced volume in the bilateral cerebellum (Table 1). *Recency*: Poor recency item performance was associated with reduced bilateral inferior cerebellar volume

#### Long Delay

*Primacy*: Poor primacy item recall at LD significantly correlated with two clusters encompassing the bilateral MTL. A left-sided cluster included large portions of the amygdala, hippocampus, entorhinal cortex, and parts of the perirhinal cortex and anterior parahippocampal cortex. This cluster also extended into the fusiform gyrus, ITG, MTG and temporopolar regions. A right-sided cluster peaked in the hippocampus and extended into the amygdala, large portions of the entorhinal and perirhinal cortices and parts of the temporal pole (Fig. 2c, Table 1). *Middle*: There were no significant clusters associated with LD recall by middle items. *Recency*: There were no significant clusters associated with LD recall by recency items.

#### Investigation of SPE change over time: Learning to Long Delay

In the next analysis, we examined whether the change in performance of primacy, middle or recency list positions from Learning to LD was associated with gray matter signal intensity (Table 1). To address this aim, we first calculated three SPE difference scores of primacy, middle, and recency positions for each participant (%-correct at LD – %-correct at Learning, Fig. 4a). We then correlated the distribution of the difference scores with gray matter volume in three separate VBM analyses. As expected, only the difference score for primacy was significantly associated with bilateral MTL integrity indicating that poorer recall of primacy items at LD relative to Learning was associated with less volume in the MTL (Fig. 4c), Table 1. This analysis revealed no significant clusters for either the recency or middle list position across Learning to LD.

**Figure 4.**
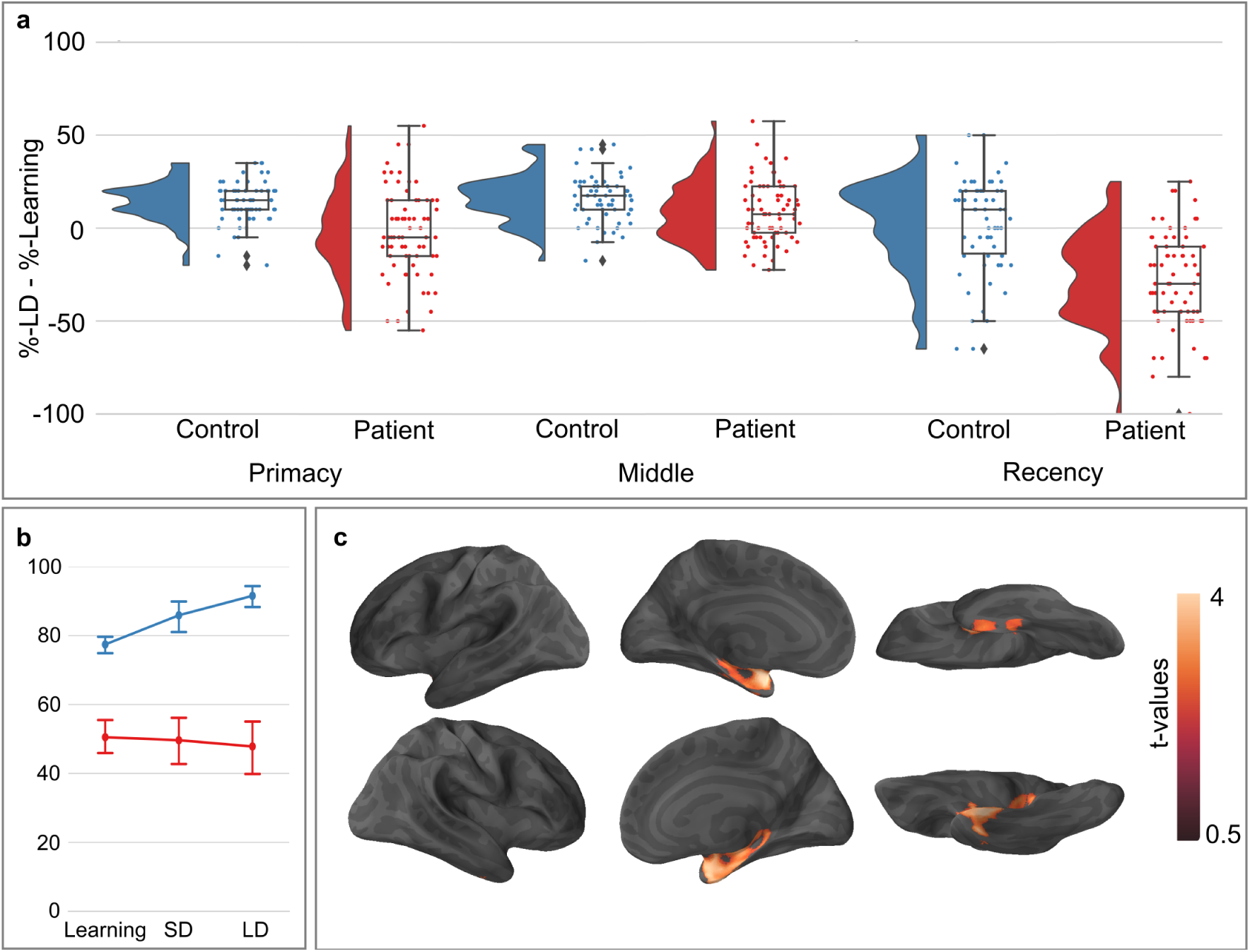
Primacy recall over time from Learning to LD: Primacy progression. (a) The raincloud plot reflects the difference scores for each serial list position, separately for healthy controls and patients. The half-violin plots on the left-hand side indicate the kernel density of the distribution of difference scores. The overlays of the jittered difference scores as a scatter plot and a boxplot are shown on the right-hand side. The box indicates the inter-quartile range (IQR), the band inside the box indicates the median, the whiskers indicate 1:5×IQR. Outliers i.e., values > 1:5×IQR are shown as diamonds. (b) The accuracy of primacy items at each time point (Learning, SD, and LD) for healthy controls (blue line) and patients (red line). Note that the primacy items in the control group significantly improve across three timepoints, i.e., ‘primacy progression’. In contrast, performance on primacy items in the patient group remains low and stable over time. (c) The correlation between gray matter volume and proportion of items recalled at long delay relative to those learned. This figure shows the regions where reduced volume is associated with poorer performance of delayed recalled relative to initial learning.

## Discussion

The present study systematically studied and found support for distinct neuroanatomical correlates for SPE at Learning, SD, and LD. We found that performance with primacy items at Learning was associated with left MTL and temporal pole volume, whereas recency item Learning was awas associated with bilateral MTL and right STG volume. Interestingly, only primacy item recall remained significantly associated with gray matter signal intensities in the lateral and medial temporal regions over time^16, 23^. Our findings further showed that healthy controls, unlike patients, improved their item recall from Learning to SD to LD. We coin this ability to improve with these items over time ‘primacy progression’. Furthermore, reduced primacy progression was associated with smaller volumes in the bilateral MTL regions. While individual SPE measures at Learning, SD, and LD were associated with distinct neuroanatomical coordinates, we newly demonstrate that the primacy progression measure may be a particularly sensitive cognitive-biomarker differentiating controls from of incipient AD-related pathology.

Early behavioral research documented that primacy item learning differentiated healthy individuals’ U-shaped from J-shaped learning curves in patients with AD^11, 13, 14, 24, 33^. The success of primacy item recall at learning reflects an ability to encode initial list items facilitated by binding of deeper semantic processing. In contrast, the J-shaped pattern found in MCI and AD characterizes the primacy impairment in semantic memory with preserved processing of recency items, reflecting reliance on the phonological rehearsal processes^3, 34^. Our corresponding VBM results demonstrated that poorer learning of primacy items was associated with reduced volume in the left ITG, MTG, temporal pole, MTL, and right inferior parietal areas. In previous studies, the temporal areas have been associated with semantic processing and memory^8, 10^. These findings add to the known temporal gyrus and hippocampal involvement associated with impaired primacy item recall found in AD^15^, amnestic MCI^35^, left hippocampal surgical excision^36^ and impaired hippocampal development^37^. The findings are also consistent with an functional MRI study demonstrating heightened BOLD activity in left MTL for primacy items probes^38^. In comparison, recency item learning was associated with reduced volume in the right STG as well as bilateral MTL. These findings reflect the importance of regions underlying auditory phonological rehearsal processes^7, 9, 30, 31^ necessary for recency item recall^3^.

Two findings differentiate the neuroanatomical correlates of primacy items of SPE over time. First at SD, primacy item recall was associated with reduced left anterolateral temporal (ITG, MTG) as well as volume in the temporopolar and right posterior temporo-occipital regions. We did not observe an association between the volume of these structures and the performance with either middle or recency items. Second, primacy item performance at LD was the only SPE measure to significantly correlate with integrity of cerebral regions. Specifically, poorer primacy item recall at LD was associated with reduced volumes of bilateral MTL, left anterolateral temporal lobe and right temporal pole. The lack of significant MTL correlation of recency items at SD or LD may reflect a floor effect given the dramatic drop in recency item performance in the patient group (see also Fig. 4a). Taken together, these findings suggest that damage to the regions, which in previous research have been shown support semantic memory processes^8, 10, 39^, disproportionately hinder primacy item recall both at SD and LD as compared to other list positions.

The poor delayed recall of recency items in the patient group supports the dual process account of instability of these items owing to the shallow, phonological rehearsal processes. While patients’ and controls’ recency item performance was comparable at learning, only patients’ recency item performance dropped dramatically by roughly half at SD with no improvement at LD. In contrast, controls’ recency item performance remained stable over time. We also showed that poorer recency item learning performance is associated with decreased bilateral MTL volume, consistent with a FDG-PET study^40^, which showed that decreased metabolic activity in MTL is associated with recency item learning. Taken together, these findings provide further evidence that while phonologically processed items may be rehearsed at learning and even SD, these shallow linguistic representations are transient and difficult for patients to consolidate (see also^41^).

Lastly, the current study also newly demonstrates that controls, but not patients, demonstrated ‘primacy progression’, which signifies that they were able to improve item recall from Learning to LD (see Fig. 1, and Fig. 2b). We propose that this measure is a critical third serial position marker, to supplement the U-versus J-shape and the recency drop. The primacy progression denotes that at Learning, items may not have been reported but were semantically encoded and hence available on later delay. This is consistent with reports of Sederberg et al.,^42^ and Schallmo et al.,^43^ who demonstrate that items activated at the time of encoding are those that are later reported. This is in stark contrast to patients with AD, whose poor initial primacy item recall at Learning represents not only the known semantic deficit^44^ and impaired consolidation^45^, but also the compounded inability to associate semantic information with the relevant ordinal position^46^. Our structural brain analyses highlight the sensitivity of primacy progression and its association with integrity of the bilateral medial temporal lobes and temporal poles consistent with the semantic and memory associations. Importantly, the remaining two contrast measures for middle and recency (LD – Learning) failed to show any significant correlations with gray matter volume.

We suggest there is a two-fold explanation for the sensitivity of primacy progression to MTL pathology in Major and Minor Neurocognitive disorder. First, this measure taps both semantic and memory processes supported by parahippocampal regions^6, 8, 27^ and second, it also reflects consolidation processes which rely heavily on the hippocampus and entorhinal cortex^8^. The neurofibrillary pathology affects these regions early in the course of Alzheimer’s disease^32^ and neurofibrillary pathology associated with cognitive decline^47^. Indeed, semantic memory impairments putatively reflecting pathology in these regions have previously been shown to be apparent well-before a clinical diagnosis of neurodegenerative disease^48^. Thus, future studies should consider the sensitivity of the primacy progression score as an early cognitive-biomarker of Alzheimer’s disease.

Word list learning is one of the most frequently used neuropsychological instruments to measure impaired episodic memory and to detect early MTL pathology in brain disease, and AD in particular. As the CVLT list-learning measure was designed to include semantic categorization, we were able to capture focal pathology reflecting the combined impoverished semantic system and verbal episodic encoding memory networks.

We note that several different SPE scoring methods exist (see comparative review,^14^), but the present study offers continued empirical validation of the sensitivity of our current regional scoring method^11, 24, 36, 49^ to regional atrophy of key neuroanatomic systems affected in neurodegenerative disease.

Limitations of the current study and future directions are worth noting. While this cross-sectional study provides a first step of cognitive-volumetric relationships of SPE profiles, future studies can delineate the longitudinal course of these measures. Future samples may also be selected to include more diverse populations, and stratify by sex. The current study addresses the SPE relationship to volume, but not to other known biomarkers associated with Alzheimer’s disease and other dementias.

The current study provides a first documentation of the functional and the neurostructural associations of primacy, middle, and recency SPE across time from initial learning to short delay followed by long delay recall. Importantly, we show that the primacy progression measure (primacy items recalled at Delay compared to primacy items correctly produced at Learning) represents a meaningful and highly sensitive cognitive-biomarker reflecting MTL pathology, which is known to be the hallmark of the development of Alzheimer’s disease. These measures provide promise for disease detection, progression, and therapeutic response.

## Methods

### Participants

Data of 236 individuals from longitudinal studies^50^ at the Basel Memory Clinic were retrieved. All individuals were over 50-years old, scored ≥ 20/30 on the Mini Mental Status Examination (MMSE)^51^ had reliable informants, and no psychiatric, neurological (other than meeting criteria for conditions below), or systemic disorders affecting cognition. Those individuals who had completed the California Verbal Learning Test (CVLT)^52^ (German version) and who underwent anatomic MRI scanning within two months (mean ± SD) = 25 ± 14 days) were selected. Healthy Control participants (HC:*n*= 62; mean age = 70.2 ± 8.3 yrs.; education = 13.2 ± 2.6 yrs.; 66 % male; mean MMSE = 29.2 ± 0.9, range 27-30) were included if they had no neurological diagnosis and if no cognitive score on the Consortium to Establish a Registry for Alzheimer’s Disease (CERAD^53^) Neuropsychological Assessment Battery fell below demographically-adjusted z-scores of −1.96 at baseline and at two-year follow-up, resulting in a ‘robust’ normative sample^54^. Patient participants (PAT: *n*= 69; mean age = 62.2 ± 9.4 yrs.; education = 13.5 ± 3.3 yrs.; 53 % male; MMSE = 28.0 ± 2.1, range 18-30) met criteria for ICD-10 criteria for Major or Minor neurocognitive disorder. Demographics of age (*t*(129) = 0.6, ns), education (*t*(129) = −0.6, ns.) and sex (*χ*^2^(1) = 2.1, ns.) were not significantly different between groups. To increase whole-brain variability of voxel-based estimates of gray matter (GM) volume, and thus validity of the brain-behavior correlations^55^, we included patients meeting criteria using classification of Major or Minor Neurocognitive disorder. The sample included Major Neurocognitive Disorder (*n* = 18)^56^; Minor Neurocognitive disorder (*n* = 46; including 44 amnestic MCI and 2 non-amnestic MCI)^57^; Lewy-Body Dementia (*n* = 3)^58^; Posterior Cortical Disorder (*n* = 1)^59^); and frontotemporal lobar degeneration (*n* = 1)^60^.

The Institutional Ethics Committees of Both Basel Cantons following principles outlined in the declaration of Helsinki and the Office of Regulatory Compliance at Queens College, City University of New York approved the study. All participants gave written informed consent prior to participation, and were reimbursed for their efforts.

### Measures

#### Regional Serial Position Measures

The CVLT^52^ 16-item word list A was divided into primacy, middle and recency list positions (4, 8, and 4 words, respectively). Serial position scores^11, 36^ were defined as items correctly recalled relative to the number presented from each list position at three time points: Learning defined as the cumulative score across Trials 1-5, SD, and LD. Note that the regional SPE scoring^11^ differs from standard CVLT-II serial position scoring^52^.

#### MRI image acquisition

High-resolution T1-weighted three-dimensional magnetization-prepared rapid acquisition gradient echo images were acquired for all participants on the same 3T MRI scanner (MAGNETOM Allegra, Siemens) using a the following acquisition parameters: TI = 1000 ms, TR = 2150 ms, TE = 3.49 ms, flip angle = 7°; rectangular field of view = 87.5 % (280 × 245 *mm*^2^), acquisition matrix = 256 × 224 mm, voxel size = 1.1 mm isotropic.

### Design and analyses

#### Behavioral analyses

Age and sex were used as a covariates in the factorial analyses to be more conservative and parallel the VBM analyses. All Accuracy scores were submitted to a mixed model analysis of covariance with group (HC, PAT) as the between-subject variable, and list position (primacy, middle, recency) and time (Learning, SD, LD) as within-subjects variables. Tukey post-hoc test significance was set at *α* = .05. The behavioral results were plotted using Python 3 (https://www.python.org/) together with Seaborn (https://github.com/mwaskom/seaborn) and the ptitprince libraries (https://github.com/pog87/PtitPrince,^61^).

#### Imaging analyses

Preprocessing and statistical analyses were conducted using SPM8 (/urlhttp://www.fil.ion.ucl.ac.uk/spm/software/) in Matlab (v2010b, Mathworks Inc.). Anatomic images were segmented into GM volumes, which were used to create a study-specific template using the DARTEL approach^62^. Individual GM images were coregistered with the study-specific template and MNI space, modulated and smoothed with an 8 mm Gaussian kernel. Each participant’s performance scores were correlated with voxel signal intensities across whole brain GM volumes using multiple regression analyses based on the General Linear Model^55, 63^. Three separate models tested the neuroanatomical correlates of primacy, middle and recency item performance at Learning, (2) SD and (3) LD. Each model included performance on all three list positions. All analyses included age, sex and total GM volume as covariates. The voxel-based correlations therefore reflect the unique relationship between performance of a single list position and regional GM volume. We report contrasts theoretically relevant to our hypotheses, namely significant positive correlations for primacy, middle and recency item performance and regional brain volumes at each time point. All analyses included total GM (to control for combined effects of brain size and overall atrophy), age and sex as covariates. For each model, the t-statistic tested for regional effects, and associated p-values were corrected for search volume with the Gaussian random field theory. Statistical parametric maps were thresholded at FDR-corrected p < .01 at cluster-level and uncorrected at the voxel level. The anatomical areas are described using the AAL atlas (^64^ and MTL subregions^65^. The t-values of significant clusters are projected onto the cortical surface using Freesurfer (https://surfer.nmr.mgh.harvard.edu/) and PySurfer (https://github.com/nipy/PySurfer/) library.

## Acknowledgements

Research in this publication was supported by Swiss National Science Foundation Ambizione Fellowship #PZ00P1_126493 to KIT, Early Postdoctoral Mobility Fellowship #BBSP_146854 and Academy of Finland grant #286070 to SLK; Professional Staff Congress–City University of New York grant #64535-00-42 and NIH-National Institute of General Medical Sciences of the National Institutes of Health (NIH-NIGMS) under award number SC3-GM122662 to NSF; MRI scan data were supported in part by research grants from GlaxoSmithKline (RES104859 – Basel) and the Novartis Foundation to AUM.

## Author contributions statement

NSF, AUM, KIT and SLK conceived the experiment; NSF, SLK and KIT performed analyses and wrote the manuscript; MS and AUM participated in patient data collecting; all authors reviewed the manuscript.

## Data availability statement

The datasets generated and analyzed in the current study contain clinical patient data. These data cannot be made publicly available due to constraints imposed by the local ethics committee at the time of the data collection.

